# The triplet codons across the watershed between the non-living and living matters

**DOI:** 10.1101/2022.09.14.508044

**Authors:** Dirson Jian Li

## Abstract

Three nucleotides per codon had been determined before deciphering the genetic code. However, it is still a mystery why there are three nucleotides per codon. This is a deceptively simple problem, which need first to clarify the prebiotic picture that has been in debate for decades. The triplet nature of life has been observed not only in the triplet codons but also in the universal 3-base periodicity in genome sequences. Here, a statistical picture on the prebiotic sequence evolution has been proposed by ascertaining the profound relationship between the evolution of the genetic code and the diversification of life. There are indications that the triplet nature of the genetic code is due to a mixture of the periods in the superhelical structures of bent DNAs.

## 1 Introduction

The triplet nature of the genetic code was first proposed by intuitively considering 4^3^ triplet codons enough for the 20 canonical amino acids before deciphering the genetic code [1] [2] [3]. It is sometimes yet mistakenly believed that the origin of the well-known three nucleotides per codon need not be further explained, just as the number 20 for canonical amino acids not. This deceptively simple problem can lead to an insight into the relationship between the evolution of the genetic code and the diversification of life (Fig 1a). There are 3 nucleotides per codon, 4 nucleotides in DNAs or RNAs, 20 canonical amino acids in proteins. How did these perplexing numbers originate and why have they remained unchanged? When choosing 3 from 4 with repetition (Fig 1b), the number of repeated permutation, namely 3^4^ = 64, agrees with the number of codons, while the number of repeated combination, namely *C*(4 + 3 − 1, 3) = 20, agrees with the number of canonical amino acids [5], which might cannot just be interpreted as a coincidence. It indicates a profound combination relationship among the above observed numbers, where the triplet nature plays a pivotal role in sequence organization. As a common feature of life, the triplet nature of life originated before the separation of the three domains of life, explanation of which leads to comprehension of the prebiotic evolution.

**Fig 1:**
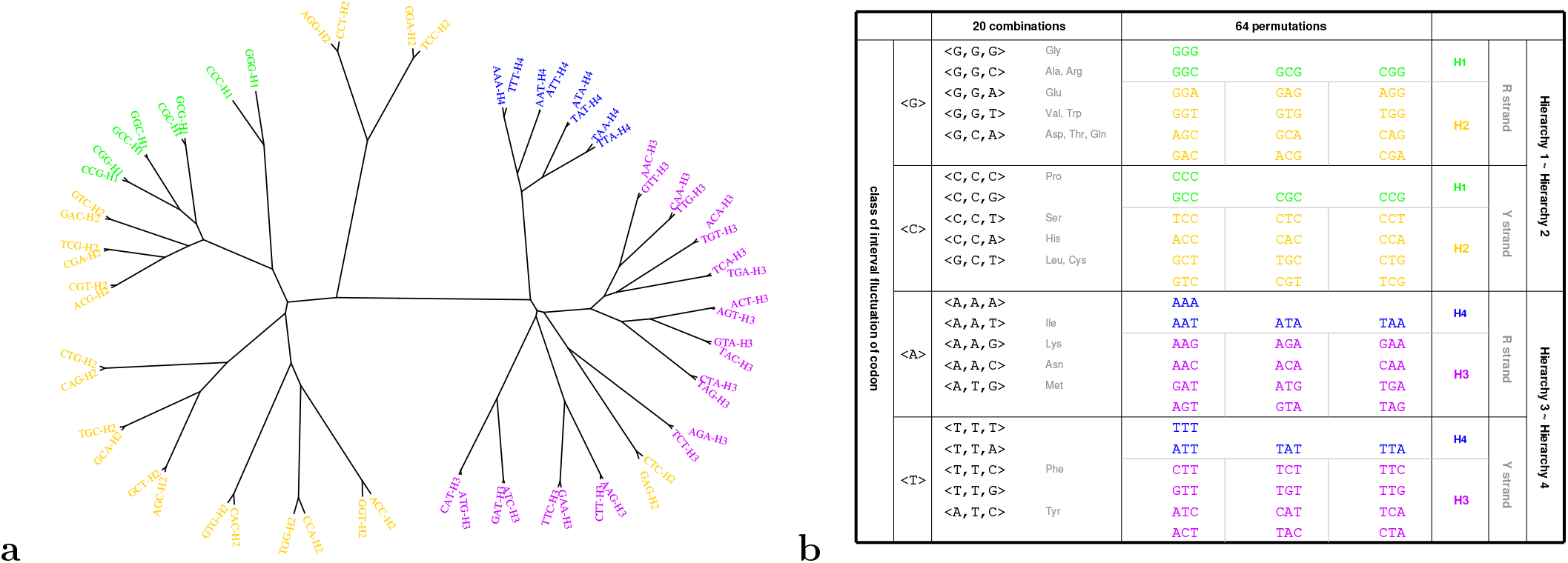
**a** The evolutionary relationship of codons by comparing codon aspect of IFCs. Both the present result on the evolution of the genetic code and the result on the diversity of life (Fig 2a in Ref. [40]) are based on the same data of IFCs. **b** The 20 groups of 64 triplet codons by the 20 repeated combinations of the 4 nucleotides, which agrees with the evolutionary Hierarchies on the roadmap (Fig 1a in Ref. [39]).

Although the origin of the genetic code has been studied for decades [6] [7] [8] [9] [10] [11] [12], the origin of the triplet nature of the genetic code has been seldom mentioned in the literature. The triplet codons was suggested to expand from doublet codons [13] [14] [15] [16] or to reduce from longer non-triplet codons [17]; it was argued that the triplet nature was forced by translocation processes with the help of fluctuations when a Brownian regime molecular machine drives the chain along the polymer in steps of three monomers [18] [19] [20]; syntheses of 4-base decoding tRNAs with improved efficiency suggested that the triplet codon code was selected because it is simpler and sufficient, not because a quadruplet codon code is unachievable [21] [22]. The theories to explain the triplet nature of the genetic code are quite different in the literature, which may weaken each other.

There was also trial to explain the 3-base periodicity in genome sequence [23]. The triplet nature has been observed not only in the translation (3 nucleotides per codon) [24] [2], but also, independently, in the neighbouring codon choice [25] [23], in the circular code [26], and in the recombination (about 3 nucleotides per RecA) [27] [28] [29] [30] [31], etc. Abnormal expansion of the triplet repeats occurred in or near genes can result in several human diseases, including myotonic dystrophy, Huntington’s disease, Spinocerebellar ataxia, Dentatorubral-pallidoluysian atrophy, etc. [32] [33]. A general theory is awaited in the literature to uniformly describe the above miscellaneous observations pertaining to the triplet nature of life all together.

The arrow of causality should be reversed to attribute the 3-base periodicity in the gene-coding region to inhomogeneous nucleotide compositions in the three codon positions [34], because this reason cannot explain the 3-base periodicity present at the level of the whole genome where the codon concept is not applicable [35]. Thereby, the correct causality direction should be that the universal 3-base periodicity in whole genome sequences determines the triplet nature of the genetic code. To avoid circular argument, the 3-base periodicity in genomes should be reinterpreted based on fundamental biophysical properties. In this paper, the 3-base periodicity in genomes has been explained by averaging the short periods in the superhelical structures of bent DNAs.

The triplet nature of the genetic code can be explained at the sequence level based on general statistical principles. After the time when RNA, DNA and protein all appeared, their sequences evolved together. Meanwhile, the prototype genomes evolved from small to large. Here, a statistical multi-period-mixing model is proposed to explain the universal 3-base periodicity in genomes by concatenating multi-periodic sequences. It has been shown that there is a significant correlation between the curvatures of bent helical DNAs and the periods of certain DNA sequences [36] [37] [38]. The universal 3-base periodicity in genome sequences has been simulated by randomly mixing and concatenating these periodic short sequences with respective periods *n* base (*n* = 2, 3, 4, 5, etc.). The universal 3-base periodicity has been observed not only in the interval fluctuation of codon (IFC) for segments whose lengths are *k* = 3 base but also in the generalised *k*-order interval fluctuation IFC^*k*^ for longer segments whose lengths are *k* base (*k* = 2, 4, 5, …) (see Methods). There is a mutual support in genome organization between these segments with different lengths *k* base (*k* = 2, 3, 4, …) for the observed periodic IFC^*k*^ and the above short sequences with multiple periods *n* base (*n* = 2, 3, 4, …) in concatenations. Such a statistical concatenation picture is able to explain the IFC^*k*^ 3-base periodicity and hence the triplet nature of the genetic code in observation.

Statistics plays an essential role in living systems at the species level, the sequence level, as well as the molecule level. Life evolves through interdependent development of these levels. During the transition from non-living to living matters, statistics played a crucial role in the origin of the genetic code and the formation of diverse genomes [39]. In the storage of evolutionary information, the triplet nature, for example three nucleotides per codon, becomes a common feature for all kinds of life. The triplet nature of life has been observed not only in the genetic code but also in the genome sequences. A 3-base periodicity has been observed in IFCs for the genomes of Bacteria, Archaea, Eukarya, and Virus [40]. Such a universal IFC 3-base periodicity at the sequence level should appear before the separation of the main branches of life, which was around the time when the triplet nature of the genetic code formed. There was, therefore, a reasonable chronological relationship between the formation of the universal IFC 3-base periodicity and the formation of the triplet nature of the genetic code.

In the previous first paper [39], the formation of the codon degeneracy can be explained according to the base substitutions during the coevolution of tRNAs with aaRSs in the prebiotic sequence evolution. In the previous second paper [40], the diversification of life can be explained according to the varying distributional features of the triplet codons in genome sequences. There is a close relationship between the above two papers, in both of which not only the triplet nature of life but also the biosynthetic families of amino acids played fundamental roles. The number of repeated combination determines the IFC types and hence the number of canonical amino acids. The results from the bottom-up study on the origin of the genetic code and the top-down study on the diversification of life agree with each other. Such an agreement increases the feasibility to clarify the origin of life by the bi-directional approach. The sequence evolution marked with the triplet nature runs through the prebiotic evolution and the consequent origin and evolution of life.

There are a few well-known common features of life, such as the genetic code, the homochirality, etc. Strictly speaking, none of them are rigorous common features. The non-standard genetic code is used differently by species [41]; the number of canonical amino acids is not always equal to 20 [42] [43]; D-amino acid residues in natural proteins perform essential biological functions [44] [45] [46]; the length of codons can deviate from 3 base according to experimental evidence on quadruplet decoding as well as the discovery of organisms with ambiguous and dual decoding [47] [48] [49] [50]. It is generally known that all matter has common physical features such as mass and spin. Although there is no rigorous common feature to distinguish living matters from non-living matters, the statistical principles at the sequence level indeed play crucial roles for the formation of all kinds of life. So life can be regarded as a kind of information system rather than a kind of matter, just as our culture differing from ourselves. The triplet nature of life plays a role of information storage format at the sequence level. Thereby, we must attach great importance to and try to explain the origin of information storage ability of life at the sequence level, based on an intensive comprehension of the picture for the statistical formation of prototype genomes.

Both the evolutionary relationship between codons and the evolutionary relationship between species have been obtained by comparing IFCs *ifc*(*int, cod, sp*) of the same genome sequences, which indicates a profound relationship between the prebiotic formation of the codon degeneracy and the subsequent diversification of life (see Methods) [39] [40]. On the one hand, the evolutionary relationship of species has been obtained by comparing IFCs *ifc*(*int, cod, sp*) for species. The three domains of life and Virus have been separated, according to biosynthetic biases of specific IFC features between different biosynthetic families of amino acids (see Fig 2a in [40]). On the other hand, the evolutionary relationship of codons (Fig 1a) has been obtained here by comparing IFCs *ifc*(*int, cod, sp*) for codons, which agrees with the four Hierarchies on the roadmap for the evolution of the genetic code (see Fig 1a in [39]). Thereby, both the information for the evolution of the genetic code and the information for the evolution of species have been stored in the statistical features of triplet codons in genome sequences. The systematic role of the triplet codons recruited at different stages for the diversification of life can be regarded as the second biological meaning of the genetic code, which is beyond its literal meaning during translation [40]. A convincing picture for the prebiotic sequence evolution is helpful to understand the pivotal role of the triplet nature of life during the formation of informative genome systems.

**Fig 2:**
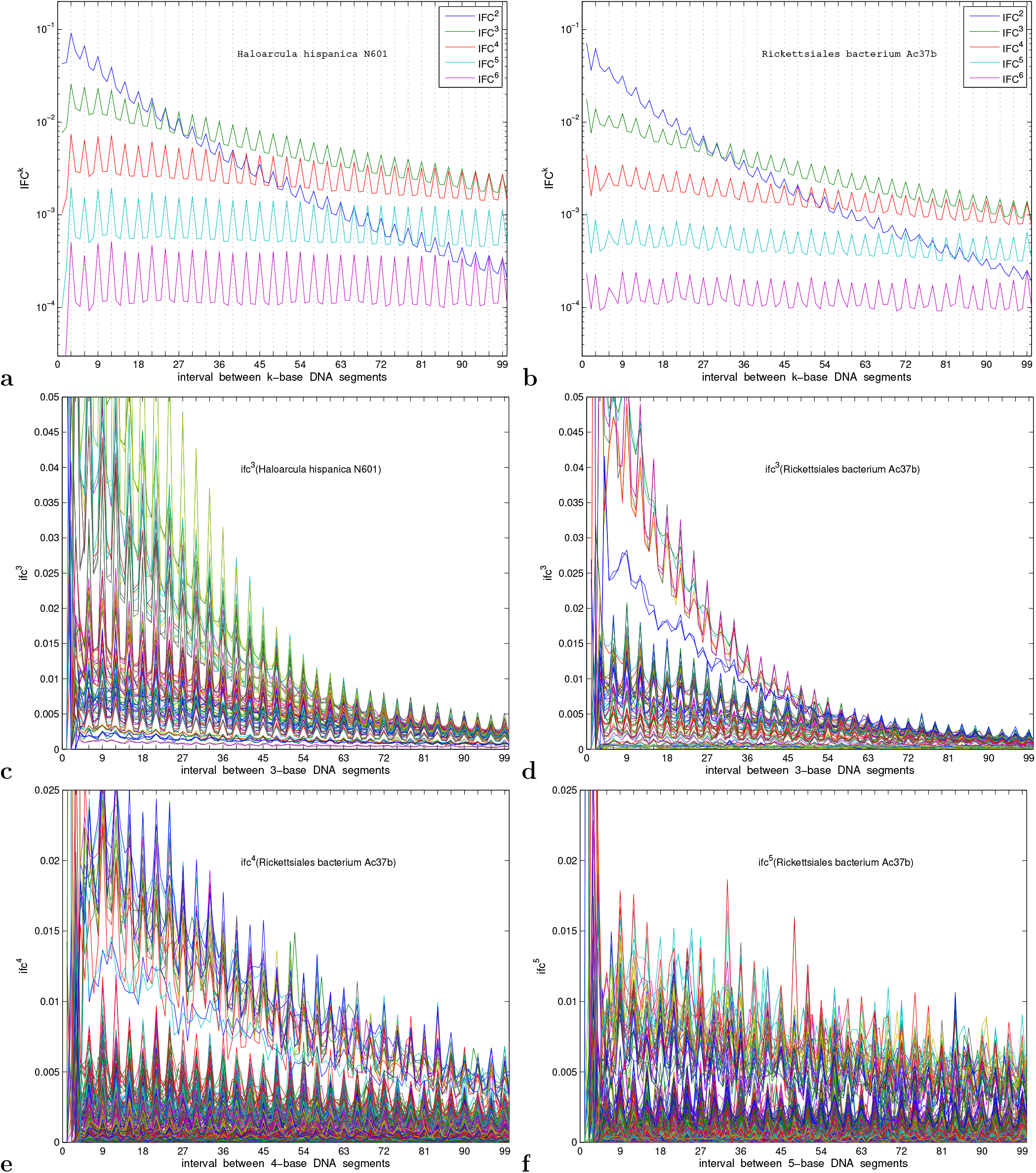
A universal 3-base periodicity for Archaea and Bacteria. **a** Haloarcula hispanica N601 (an archaeon). **b** Rickettsiales bacterium Ac37b (a bacterium). **c** *ifc*^3^ of Haloarcula hispanica N601. **d** *ifc*^3^ of Rickettsiales bacterium Ac37b. **e** *ifc*^4^ of Rickettsiales bacterium Ac37b. **f** *ifc*^5^ of Rickettsiales bacterium Ac37b.

## Results

### Relationship between the evolution of the genetic code and the diversification of life

There are a couple of common features between before and after the origin of life, when comparing the formation of the codon degeneracy [39] and the diversification of life [40]. First, the triplet nature of both the genetic code and IFCs is an intrinsic nature of life; second, biosynthetic families of amino acids help both the expansion of the genetic code and the diversification of life [39] [40]. Both the evolutionary relationship of the triplet codons (Fig 1a) and the evolutionary relationship of species (Fig 1d, 2a in [40]) can be obtained by comparing the statistical features of codons in genomes of same group of species (see Methods). Such features across the transition from non-living and living help us to understand the origin of life from both bottom-up approach and top-down approach.

#### (1) The triplet nature of life across the watershed between the non-living and living matters

There are two interrelated manifestations for the triplet nature of life: the well-known triplet codons, and the universal IFC 3-base periodicity [40]. The triplet codons should appear before the origin of life, while the universal 3-base periodicity has been observed unexceptionally in IFCs of all kinds of contemporary species (Fig 2, 3) [40]. The triplet nature of life at the sequence level has remained across the prebiotic and the consequent origin and evolution of life, and becomes a near-rigorous common feature for all kinds of life. IFC *ifc*(*int, cod, sp*) and reduced IFC *IFC*(*int, sp*) can be generalised to *k*-order IFC *ifc*^*k*^(*int, cod, sp*) and *k*-order reduced IFC *IFC*^*k*^(*int, sp*), in all of which the universal 3-base periodicity has been observed for Bacteria, Archaea, Eukarya, and Virus (Fig 2, 3, and Fig S1-S8)(see Methods). Protein length distributions in observation indicate that long proteins can be concatenated by short segments [51]. According to the statistical multi-period-mixing model (see Methods), the universal 3 base period is due to the average of different periods of the bent segments of the prototype genome. The sequence evolution led to the triplet nature of life, as a standard genome format.

**Fig 3:**
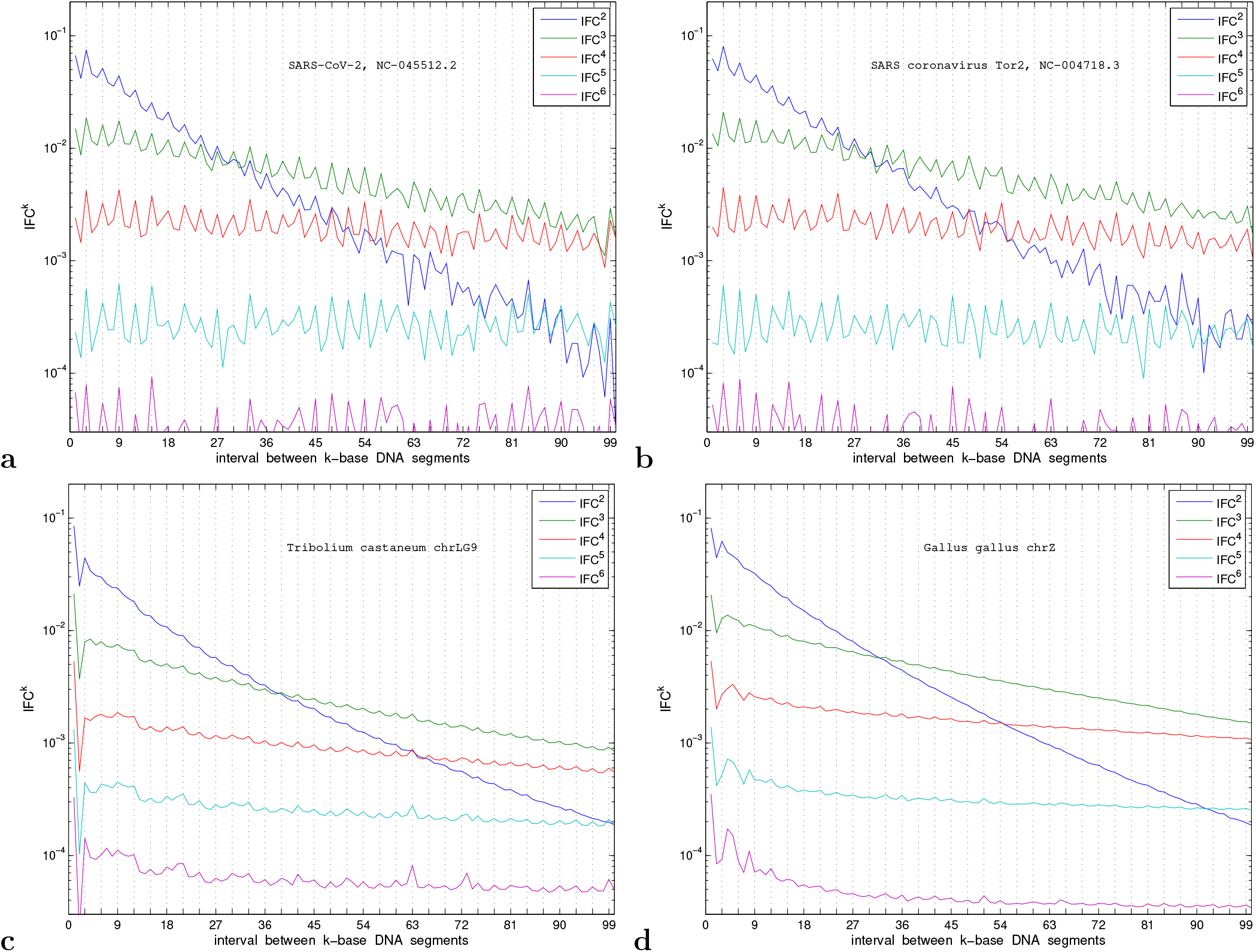
A universal 3-base periodicity for Virus and Eukarya. **a** SARS-CoV-2. **b** SARS coronavirus Tor2. **c** Tribolium castaneum chrLG9 (an insect). **d** Gallus gallus chrZ (a bird).

The formation of the triplet nature of the genetic code can be explained based on the formation of prototype genomes by concatenating short periodic sequences during phased evolution of the genetic code (see Methods). Circular code has been found as an ancestral structural property of genes to retrieve the reading frame locally [26] [52] [53] [54] [55] [56]. The 12 amino acids encoded by the 20 codons in the circular code *X*_0_ are as follows: *Gly, Ala, Glu, Asp, V al, Leu, Thr, Ile, Phe, Asn, Gln*, and *Tyr* [52], the former seven of which are encoded by codons in the initial subset on the roadmap (Fig 1a in [39]), and the next three of which are situated next to the initial subset on the roadmap. The common amino acids encoded early by the codons in both the circular code *X*_0_ and the initial subset on the roadmap belong to Phase *I* amino acids [57] [58]. The amplitude of fluctuations in IFC is an evolutionary feature, which relates to the subset of codons and the corresponding amino acids. The IFC amplitudes become high when concatenating circular code in generation of the periodic segments in the model (see Fig 4a in [40]), which accounts for the high IFC amplitudes of Bacteria and Archaea that appeared early before the fulfillment of the genetic code. According to the simulation, additionally more number of codons in the proper subset of the 64 triplet codons brings about low IFC amplitudes, which corresponds to Eukarya that appeared late. Except for the universal IFC 3-base periodicity, there are specific IFC features for different species, such as the varying amplitudes of the fluctuations, the varying heights of distributions for different codons, and the steep peaks in the IFCs especially for eukaryotes with more non-coding DNAs (Fig S1-S8). It has also been observed that the 3-base-period structure appeared mainly in the coding regions in genomes [25] [59] [34] [60]. The validity of the statistical multi-period-mixing model can be confirmed by the proper simulations of complicated specific IFC features (Fig 4).

**Fig 4:**
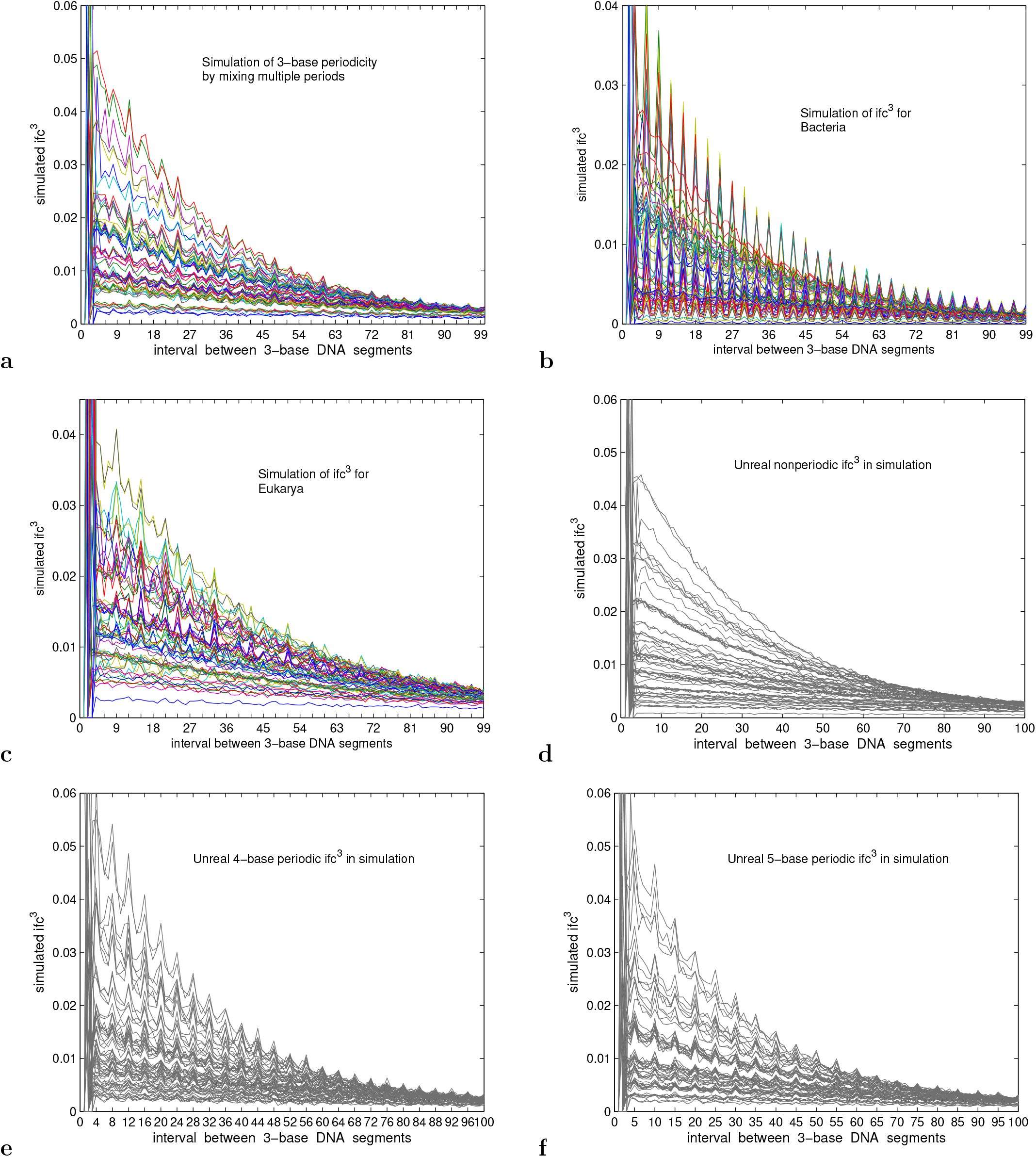
Simulation of the universal 3-base periodicity. **a** Simulation of 3-base periodicity by mixing multi-periodic sequences. **b** Simulation of *ifc*^3^ for Bacteria. **c** Simulation of *ifc*^3^ for Eukarya. **d** Unreal smooth distribution by simulation. **e** Unreal 4-base periodicity by simulation. **f** Unreal 5-base periodicity by simulation.

#### (2) Biosynthetic families of amino acids across the watershed between the non-living and living matters

The biosynthetic families of amino acids play crucial roles in both the expansion of the genetic code during coevolution of tRNAs with aaRSs and the diversification of life, which facilitated the prebiotic and the consequent evolution of life. During the intricate coevolution of the aaRSs and tRNAs and the cooperative recruitment of codons and amino acids (Fig 7, 3a in [39]), the biosynthetic families of amino acids (*Glu* family, *Asp* family, *V al* (pyruvate) family, *Ser* family, and *Phe* (aromatic) family) helped the recruitments of the late amino acids alongside the early amino acids *Ala, Pro, Ser, Thr, V al*, and *Leu* in the initial subset on the roadmap (Fig 7, 3a, 1a in [39]). There is rich evolutionary information in the specific IFC features of different codons (Fig 2, 3). The three domains of life and Virus have been separated in the 3-dimensional biosynthetic bias space whose coordinates are *Ser*−*Val* bias, *Phe* − *Ser* bias, and *Asp* − *Glu* bias, respectively, owing to the evolutionary aspect of the genetic code on the different distributional features among codons recruited at different stages (Fig 2 in [40]).

### Explanation of the triplet nature in the genetic code and in the genome sequences

It is feasible to explain the 3-base periodicity by the correlation between the superhelical structures of bent DNAs and the repeated sequences based on the following observations. There is a close relationship between DNA sequences and DNA structural properties beyond double helix [61]. It was also suggested that the 3-base periodicity for neighboring codons in DNA sequences may be associated to local RNA secondary structure [62]. Bent DNAs result when special base sequences or structural motifs are repeated in phase with the DNA helical repeat [63]. Evidence was presented for periodic repetition of particular dinucleotides as facilitators of bending [51]. Bending of DNA results in diminution of its total winding angle, this fact probably being important for the supercoiled duplexes [64]. It is suggested that nucleotide composition has a widespread and direct influence on chromatin structure [65]. Although a whole genome level understanding of the correlation between eukaryotic chromatin structures and the features of genome sequences is still limited [66] [67], the correlation between local DNA structures and short special base sequences eventually results in the correlation between global chromatin structures and whole genome specific sequences.

The bases are situated alternatively at the inner ring and the outer ring along a superhelical structure of bent DNA. It has been observed that AT-rich sequences tend to appear in the inner ring of nucleosome than in the outer ring of nucleosome [68] [63]. There are regular bent DNAs around histones for Eukarya and histone like proteins for Bacteria and Archaea. The alternative circumstances around the bases bring about the periodic sequence along the bent DNA with a certain curvature. Thereby, regular superhelical structure of a bent DNA may correspond to a periodic sequence. In addition, there are numerous segment periods that correspond to different curvatures of segments of a freely bent DNA. Thus, the mixture of different segment periods leads to the 3-base periodicity in the IFC of the whole sequence (see Methods). The genes can originate from ancestrally nongenic sequences [69] [70]. The appearance of triplet codons can improve the efficiency to form peptide bonds in prebiotic evolution. Even if a prototype ribosome may not assist to align the amino acids, the triplet codons can provide respective three interaction points along the mRNA so as to align the carried amino acids before forming peptide bonds by fixing the orientation angles of tRNAs.

The universal IFC 3-base periodicity plays a role of genome format in storage of biological evolutionary information, just like the role of the file format on computer data storage system. When concatenating numerous segments into a genes for the earliest informative macromolecules, such a 3-base genome format makes the pace neat so that the biological functions of the segments faithfully transfer to the functions of the genes. The biological genes were selected efficiently by the criteria of the triplet nature, when candidate segments are situated, with the greatest probability, by every three bases. The biological evolutionary information is stored in the same 3-base genome format, no matter how these meaningful segments are combined randomly. Such a 3-base genome format ensures that same codons are situated, with the greatest probability, by every three bases, which is able to improve the efficiency of the match between the codons and the corresponding tRNAs. The efficiency can be maintained when the number of canonical amino acids equals to the 20 types of IFCs according to the repeated combination of 4 nucleotides with specific compositions. There should be a good agreement between the frequencies of the 64 codons and the abundances of the respective tRNAs with complementary anti-codons, violation of which would result in fatally low translation efficiency.

## Discussions and Conclusions

The transition from non-living to living matters needs the mutual benefit between the prebiotic chemical evolution and the macromolecular conformation evolution. We might never experimentally achieve the spontaneous emergence of a living system absolutely from the non-living matters, because this process with numerous uncertain events has to spend considerable time much longer than the whole history of human beings. What we can do is to study the relationship between the evolution before and after the origin of life so as to approach the truth of this event bi-directionally. The aim of this paper is to attain a reasonable picture to explain the triplet nature of the genetic code. A rough sketch has been provided here for the early history of life, where the statistical principles in the sequence evolution are cherished. The simulation of the observed 3-base IFC periodicity shows that the triplet nature of the genetic code is due to the mixture of the intrinsic periods in the superhelical structures of bent DNAs. The triplet nature at the sequence level remains as a pivotal feature of life across the watershed between the non-living and living matters, which underlies both the evolution of the genetic code and the diversification of life.

The prebiotic picture remains in debate and should be clarified. Pragmatically, we can compare the evolution before and after the origin of life, then the picture can be inferred bi-directionally from the same or quite similar properties between them. Concretely speaking, the triplet nature of the genetic code formed before the origin of life, and the IFC 3-base periodicity remains unchanged after the origin of life. The triplet nature of life, as a standard storage format, facilitated efficiency during the transition from non-living to living matters. Furthermore, the biosynthetic families of amino acids facilitated both the expansion of the genetic code and the formation of distributional features of triplet codons for the variety of life. Both the triplet nature of life and the biosynthetic families of amino acids contributed to the common and specific features in sequence evolution across the watershed between the non-living and living matters. The universal IFC 3-base periodicity played a role of data storage format at the sequence level, while the biosynthetic families of amino acids brought about different sequence combinations. Numerous genomes with same format and different combinations emerged and evolved based on statistical principles at the sequence level. The present bi-directional approach helps clarify the picture for the prebiotic evolution.

## Methods

### Interval fluctuation of codon and its generalisation

The interval fluctuation of codon (IFC for short) is defined by the distribution of codon intervals in the genome sequence of a species, where rich evolutionary information is stored in both the universal IFC 3-base periodicity and the specific IFC features. An IFC *ifc*(*int, cod, sp*) consists of 64 normalized distributions of intervals between the respective triplet codons *cod* in the complete genome sequence of a certain species *sp*. Concretely speaking, let the 3-base segment *b*′*b*′′*b*′′′ correspond to any triplet codon, where *b*′, *b*′′ and *b*′′′ are one of the four nucleobases *G, C, A*, and *T*, respectively. And let the sequence *s*_1_*s*_2_…*s*_*i*−1_*s*_*i*_*s*_*i*+1_…*s*_*j−*1_*s*_*j*_*s*_*j*+1_…*s*_*z*−1_*s*_*z*_ represent the complete genome sequence of a species *sp* whose genome size is *z*. The complete genome sequences in the calculations have been downloaded from NCBI. A codon interval is defined as the difference *j* − *i* for a certain *cod* = *b*′*b*′′*b*′′′, if and only if both *s*_*i*_*s*_*i*+1_*s*_*i*+2_ and *s*_*j*_*s*_*j*+1_*s*_*j*_+2 are the pair of neighbouring same *b*′*b*′′*b*′′′. These intervals play essential roles in genome organization. A series of codon intervals are obtained for *cod* = *b*′*b*′′*b*′′′ after scanning the above complete genome sequence from left to right. Let *ifc*_0_(*int, cod, sp*) = *m* denotes the distribution of codon intervals for *cod* if there are totally *m* codon intervals in the above series of codon intervals whose values are equal to *int*, where *int* = 1, 2, …, *ct*, and the cutoff *ct* = 1000. The IFC is the normalized distribution of codon *cod* for species *sp*, namely

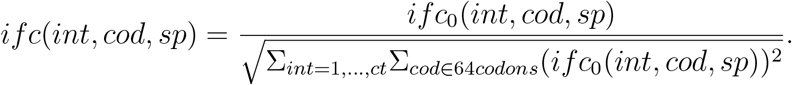

The influence of the great variation in genome sizes has been eliminated by the normalization in the definition. Then, the reduced IFC for species *sp* is defined as the mean of the 64 normalized distributions *ifc*(*int, cod, sp*) as follows:

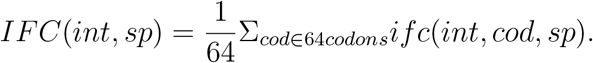

Furthermore, the above definition can be generalised to the *k*-order IFC *ifc*^*k*^(*int, cod, sp*) for species *sp* as follows, where the universal 3-base periodicity can also be observed (Fig 2, 3). If replacing the 3-base segment *b*′*b*′′*b*′′′ by the *k*-base segment *b*_1_*b*_2_…*b*_*k*_ (*k* = 2, 3, 4, 5, …), similarly, the *k*-order IFC is defined by the set of 4^*k*^ normalized *k*-order distributions *ifc*^*k*^(*int, b*_1_*b*_2_…*b*_*k*_, *sp*) (*k* = 2, 3, 4, 5, …, *k* can be omitted when it is 3). And, the *k*-order reduced IFC is defined by the mean of 4^*k*^ normalized *k*-order distributions *ifc*^*k*^(*t, b*_1_*b*_2_…*b*_*k*_, *sp*) as follows:

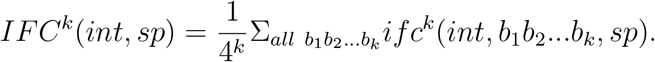

For a species with several chromosomes, these sequences can be concatenated into a whole sequence. Note that the concatenation order of the chromosomes does not influence the result. The IFC *ifc*(*int, cod, sp chr*) for a species *sp* can also be defined for each chromosome *chr*. Interestingly, the patterns of IFC and the generalised *k*-order IFC of a certain species are almost the same for different chromosomes. The evolutionary information is stored in the IFCs and the generalised *k*-order IFCs, which are generally similar for the closely related species.

### Obtaining evolutionary relationships for both codons and species by comparing IFCs

When comparing the codon aspect of IFCs, the IFC correlation coefficient for species *sp* between a pair of codons *cod*_1_ and *cod*_2_ is defined as follows:

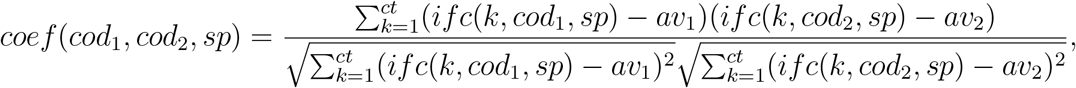

Where 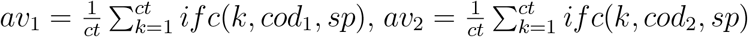. And the IFC correlation coefficient *Coef* (*cod*_1_, *cod*_2_) between a pair of codons *cod*_1_ and *cod*_2_ is defined as follows: 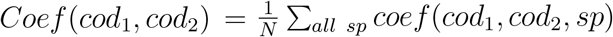. The distance matrix for the 64 codons is defined as *M*_*codon*_(*cod*_*i*_, *cod*_*j*_) = 1 − (*Coef* (*cod*_*i*_, *cod*_*j*_) + 1)*/*2, based on which an evolutionary relationship of the 64 codons (Fig 1a) has been constructed by PHYLIP software [71] [72].

When comparing the species aspect of IFCs, the three domains of life and Virus have been separated by comparing IFCs with consideration of the biosynthetic biases between groups of codons corresponding to different biosynthetic families of amino acids (Fig 2a in Ref. [40]). In both of the above calculations, 952 complete genomes (748 bacteria, 55 archaea, 16 eukaryotes and 133 viruses) downloaded from NCBI have been used, without loss of generality.

### Simulation of the IFC 3-base periodicity based on a statistical multi-period-mixing model

The 3-base periodicity observed in IFCs for all kinds of life has been simulated in a multi-period-mixing model for early genome assembly. In the simulation, some short periodic sequences have been generated, considering the correspondence between the superhelical structures of bent DNAs and their repeated sequences [61] [63]. Then, the genome can be assembled by randomly concatenating the short periodic sequences. The curvatures of segments of genome of a species vary in a certain range, which result in the corresponding range of periods of the segments. In simulation, the variation range of periods of the segments is set from 2 base to 9 base. The prototype genome sequences have been obtained by randomly concatenating short sequences with different periods by the statistical multi-period-mixing model. Amazingly, without loss of generality, numerous simulations show that the base is the most obtained period in the simulated whole genomes by mixing sequences with different periods (Fig 4a). The reason is that 3 is a usual factor of the small numbers. And, 3 is also the average of the multiplication factors of numbers from 2 to 9, or the approximate average of the multiplication factors of numbers from 2 to a not too large number.

The phased evolution of the genetic code has been considered in generation of the periodic sequences by the statistical multi-period-mixing model [40]. Evidences show that some amino acids were recruited earlier than others [57] [58], thereby, only codons in a proper subset of the 64 codons encode the early recruited amino acids. When generating 3-base periodicity, for each time, some triplet codons are chosen randomly from a certain proper subset of the 64 codons, when concatenating them into a short sequence. Different short sequences can be obtained from different proper subsets that may have evolved in different environments. The prototype genome sequence was obtained by concatenating these short periodic sequences, each of which were obtained by concatenating codons from respective proper subsets. Thus, the 3-base periodicity has been observed in the IFCs of the simulated genomes (Fig 4b). The number of elements in the proper subset influences the amplitude of fluctuations in the simulated IFC. More number brings about low fluctuations. If choosing from the complete set of the 64 codons, no periodic fluctuations have been observed in the simulated IFC (Fig 4d). This can be generalised for generating k-base periodicity. If choosing from proper subsets of the 4^4^ quadruplet permutations of the four nucleotides, 4-base IFC periodicity can be easily simulated (Fig 4e). If choosing from proper subsets of the 4^5^ quintuplet permutations of the four nucleotides, 5-base IFC periodicity can be easily simulated (Fig 4f). The proper subsets can change randomly, but the k-base periodicity remains unchanged if choosing from the same k-base permutation proper subsets. Integrating the above results, numerous periodic sequences with randomly different periods from 2 base to 9 base have been generated, with the same weight, by the statistical multi-period-mixing model, then the prototype genomes can be obtained by concatenating these short sequences with random periods. Thus, the 3-base period has been obtained in the IFCs of these simulated prototype genomes (Fig 4a), which shows that the IFC 3-base periodicity comes statistically from the average effect of these different periods.

Furthermore, the specific IFC features for different species, especially the complicated specific IFC features for eukaryotes, can also be simulated, which confirms the validity of the picture of the formation of prototype genomes in the statistical multi-period-mixing model. Except for the IFC 3-base periodicity, there are specific IFC features for different species, such as the varying amplitudes of the fluctuations, the varying heights of distributions for different codons, and the steep peaks in the IFCs especially for eukaryotes with more non-coding DNAs. If choosing from the proper subsets with less elements at early stage or more elements at late stage of the permutations, the amplitudes of the IFC fluctuations become high or low in the simulations, which accounts for the high amplitudes for early Bacteria and Archaea and low amplitudes for late Eukarya. If choosing more GC-rich codons for simulated GC rich genome, the distributions corresponding to GC-rich codons are higher than the distributions corresponding to AT-rich codons, which account for the varying heights of distributions between different codons for genomes with different GC contents. If repeatedly concatenating identical codons or longer segments, or alternatively and repeatedly choosing from two proper subsets of the 64 codons, the observed complex steep peaks and alternative periods in IFCs of eukaryotes can be simulated (Fig 4c). The powerful ability to simulate the complicated IFC features of genome of different species is due to the convincing statistical picture for the formation of prototype genomes in the multi-period-mixing model.

## Supporting Information

The supplementary figures Fig S1-S8 can be found online.

## Acknowledgments

This study was funded by the Fundamental Research Funds for the Central Universities. Thanks go to Jinyi Li for valuable discussions.

## References

[1] Gamow, G. (1957) Voprosy Biofiziki (Problems in Biophysics). Moscow: Inostr. Lit. p. 203.

[2] Crick, F. H. C., Barnett, L., et al. (1961) General nature of the genetic code for proteins. Nature 192:1227–1232.

[3] Nirenberg, M. W., Matthaei, J. H. (1961) The dependence of cell-free protein synthesis in E. coli upon naturally occurring or synthetic polyribonucleotides. Proc. Natl. Acad. Sci. U. S. A. 47(10):1588–1602.

[4] Gamow, G., Matter, Earth, and Sky (Englewood Cliffs, Prentice-Hall, 1965).

[5] Chernin, A. D. (1994) Gamow in America: 1934-1968 (On the ninetieth anniversary of G A Gamov’s birth). Physics-Uspekhi 37:791–802.

[6] Gonzalez, D. L., Giannerini, S., Rosa, R. (2019) On the origin of degeneracy in the genetic code. Interface Focus 9:20190038.

[7] Sengupta, S., Higgs, P. G. (2015) Pathways of Genetic Code Evolution in Ancient and Modern Organisms. J. Mol. Evol. 80:229–243.

[8] Knight, R. D., Freeland, S. J., Landweber, L. F. (2001) Rewiring the keyboard: evolvability of the genetic code. Nat. Rev. Genet. 2:49–58.

[9] Wong, J. T. (1975) A coevolution theory of the genetic code. Proc. Natl. Acad. Sci. U. S. A. 72:1909–1912.

[10] Klipcan, L., Safro, M. (2004) Amino acid biogenesis, evolution of the genetic code and aminoacyl-tRNA synthetases. J. Theor. Biol. 228(3):389–396.

[11] de Pouplana, L. R., ed., The Genetic Code and the Origin of Life (Kluwer Academic, New York, 2004).

[12] Crick, F. H. C. (1968) The origin of the genetic code. J. Mol. Biol. 38:367–379.

[13] Jukes, T. H. (1973) Possibilities for the evolution of the genetic code from a preceding form. Nature 246:22–26.

[14] Wilhelm, T., Nikolajewa, S. (2004) A new classification scheme of the genetic code. J Mol Evol. 59:598–605.

[15] Wu, H. L., Bagby, S., Jean, M. H., Elsen, D. V. (2005) Evolution of the genetic triplet code via two types of doublet codons. J. Mol. Evol. 61:54–64.

[16] Alejandro, F., Tom, F. (2018) The standard genetic code can evolve from a two-letter GC code without information loss or costly reassignments. Orig. Life Evol. Biosph. 48:259–272.

[17] Baranov, P. V., Venin, M., Provan, G. (2009) Codon size reduction as the origin of the triplet genetic code. Plos One 4:e5708.

[18] Martínez-Mekler, G., Cázarez-Bush, F., Aldana, M. et al. (1996) Origin of the three base codon composition: A Brownian regime molecular machine approach. Orig. Life Evol. Biosph. 26:311.

[19] Martínez-Mekler, G., Aldana, M., Cázarez-Bush, F., García-Pelayo, R., Cocho, G. (1999) Primitive molecular machine scenario for the origin of the three base codon composition. Orig. Life Evol. Biosph. 29:203–214.

[20] Aldana-González, M., Cocho, G., Larralde, H., Martínez-Mekler, G. (2003) Translocation properties of primitive molecular machines and their relevance to the structure of the genetic code. J. Theor. Biol. 220:27–45.

[21] DeBenedictis, E. A., Söll, D., Esvelt, K. M. (2022) Measuring the tolerance of the genetic code to altered codon size. eLife 11:e76941.

[22] de la Torre, D., Chin, J. W. (2021) Reprogramming the genetic code. Nature Reviews. Genetics 22:169–184.

[23] Sánchez, J. (2011) 3-base periodicity in coding DNA is affected by intercodon dinucleotides. Bioinformation 6(9):327–329.

[24] Dounce, A. L. (1952) Duplicating mechanism for peptide chain and nucleic acid synthesis. Enzymologia 15(5):251–258.

[25] Li, W. (1997) The study of correlation structures of DNA sequences: a critical review. Computers Chem. 21:257–271.

[26] Arqués, D. G., Lapayre, J.-C., and Michel, C. J. (1995) Identification and simulation of shifted periodicities common to protein coding genes of eukaryotes, prokaryotes and viruses. J. Theor. Biol. 172:279–291.

[27] Chen, Z., Yang, H., Pavletich, N. P. (2008) Mechanism of homologous recombination from the RecA-ssDNA/dsDNA structures. Nature 453:489–494.

[28] Eriksson, S., Nordén, B., Takahashi, M. (1990) Binding of DNA quenches tyrosine fluorescence of RecA without energy transfer to DNA bases. J. Biol. Chem. 268(3):1805–10.

[29] Bell, C. E. (2005) Structure and mechanism of Escherichia coli RecA ATPase. Mol. Microbiol. 58:358–366.

[30] Capua, E. D., Engel, A., Stasiak, A., Koller, T. (1982) Characterization of complexes between RecA protein and duplex DNA by electron microscopy. J. Mol. Biol. 157(1):87–103.

[31] Egelman, E., Yu, X. (1989) The location of DNA in RecA-DNA helical filaments. Science 245:404–407.

[32] Wang, Y., Amirhaeri, S., Kang, S., Wells, R., Griffith, J. (1994) Preferential nucleosome assembly at DNA triplet repeats from the myotonic dystrophy gene. Science, 265(5172), 669-671.

[33] Sinden, R. R., Potaman, V. N., Oussatcheva, E. A., Pearson, C. E., Shlyakhtenko, L. L. S. (2002) Triplet repeat DNA structures and human genetic disease: dynamic mutations from dynamic DNA. Journal of Biosciences 27:53–65.

[34] Yin, C., Yau, S. S.-T. (2005) A Fourier characteristic of coding sequences: origins and a non-Fourier approximation. J. Comp. Biol. 12:1153–1165.

[35] Shah, K., Krishnamachar, A. (2012) On the origin of three base periodicity in genomes. Biosystems 107:142–144.

[36] Moreno-Herrero, F., Seidel, R., Johnson, S. M., et al. (2006) Structural analysis of hyperperiodic DNA from Caenorhabditis elegans. Nucl. Acids Res. 34:3075–3066.

[37] Rieko, O., Maiko, H., Michio, O., Ryoiti, K. (1998) Conservation and continuity of periodic bent DNA in genomic rearrangements between the c-myc and immunoglobulin heavy chain mu loci. Nucl. Acids Res. 26:3026–33.

[38] Zhou, K., Aertsen, A. Michiels, C. W. (2014) The role of variable DNA tandem repeats in bacterial adaptation. FEMS Microbiol. Rev. 38:119–141.

[39] Li, D. J. (2021) Formation of the codon degeneracy during interdependent development between metabolism and replication. Genes 12: 2023.

[40] Li, D. J. (2022) Distributional features of triplet codons in genomes underlie the diversification of life. Biosystems 217:104681.

[41] Santos, M., Ueda, T., Watanabe, K., Tuite, M. F. (2010) The non-standard genetic code of candida spp.: an evolving genetic code or a novel mechanism for adaptation? Molecular Microbiology 26:423–431.

[42] Prat, L., Heinemann, I. U., et al. (2012) Carbon source-dependent expansion of the genetic code in bacteria. Proc. Natl. Acad. Sci. U. S. A. 109:21070–21075.

[43] Zhang, Y., Baranov, P. V., Atkins, J. F., Gladyshev, V. N. (2005) Pyrrolysine and selenocysteine use dissimilar decoding strategies. The J. Biol. Chem. 280:20740–51.

[44] Mitchell, J. B. O., Smith, J. (2003) D-amino acid residues in peptides and proteins. Proteins: Structure, Function, and Bioinformatics 50(4):563–571.

[45] Heck, S. D., Siok, C. J., et al. (1994) Functional consequences of posttranslational isomerization of Ser(46) in a calcium-channel toxin. Science 266:1065–1068.

[46] Kreil, G. (1994) Conversion of L-to D-amino acids: a posttranslational reaction. Science 266:996–997.

[47] Klobutcher, L. A., Farabaugh, P. J. (2002) Shifty ciliates: frequent programmed translational frameshifting in euplotids. Cell 111:763–766.

[48] Gomes, A. C., Miranda, I., Silva, R. M., Moura, G. R., Thomas, B., et al. (2007) A genetic code alteration generates a proteome of high diversity in the human pathogen Candida albicans. Genome Biol. 8:R206.

[49] Tamas, I., Wernegreen, J. J., Nystedt, B., Kauppinen, S. N., Darby, A. C., et al. (2008) Endosymbiont gene functions impaired and rescued by polymerase infidelity at poly(A) tracts. Proc. Natl. Acad. Sci. U. S. A. 105:14934–14939.

[50] Turanov, A. A., Lobanov, A. V., Fomenko, D. E., Morrison, H. G., Sogin, M. L., et al. (2009) Genetic code supports targeted insertion of two amino acids by one codon. Science 323:259–261.

[51] Trifonov, E. N., Sussman, J. L. (1980) The pitch of chromatin DNA is reflected in its nucleotide sequence. Proc. Natl. Acad. Sci. U. S. A. 77:3816–3820.

[52] Arqués D. G., and Michel C. J. (1996) A complementary circular code in the protein coding genes. J. Theor. Biol. 182:45–58.

[53] Xu, L., Liu J. K., Wong, T. Y. (2012) Bacterial phylogenetic tree construction based on genomic translation stop signals. Microb. Inf. Exp. 2, 6.

[54] Seligmann, H. (2018) Alignment-based and alignment-free methods converge with experimental data on amino acids coded by stop codons at split between nuclear and mitochondrial genetic codes. BioSystems 167:33–46.

[55] Ahmed, A., Frey G., and Michel C. J. (2010) Essential molecular functions associated with the circular code evolution. J. Theor. Biol. 264:613–622.

[56] Michel, C. J. (2017) The Maximal C3 Self-complementary trinucleotide circular code X in genes of Bacteria, Archaea, Eukaryotes, Plasmids and Viruses. Life 7, 20.

[57] Wong, J. T. (1981) Coevolution of the genetic code and amino acid biosynthesis. Trends Biochem. Sci. 6:33–36.

[58] Pizzarello, S., Meteorites and the chemistry that preceded life’s origin. in Wong, J.T.; Lazcano, A., Ed., Prebiotic Evolution and Astrobiology (Landes Bioscience: Austin, TX, USA, 2009).

[59] Fickett, J. W., Tung, C.-S. (1992) Assessment of protein coding measure. Nucl. Acid. Res. 20:6441–6450.

[60] Yin, C., Yau, S. T. (2007) Prediction of protein coding regions by the 3-base periodicity analysis of a DNA sequence. J. Theor. Biol. 247:687–694.

[61] Baldi, P., Baisnée, P.-F. (2000) Sequence analysis by additive scales: DNA structure for sequences and repeats of all lengths. Bioinformatics 16:865–889.

[62] Duan, J., Antezana, M. A. (2003) Mammalian mutation pressure, synonymous codon choice, and mRNA degradation. J. Mol. Evol. 57:694–701.

[63] Crothers, D. M., Haran, T. E., Nadeau, J. G. (1990) Intrinsically bent DNA. Journal of Biological Chemistry, 265(13):7093–6.

[64] Zhurkin, V. B., Lysov, Y. P., Ivanov, V. I. (1979) Anisotropic flexibility of DNA and the nucleosomal structure. Nucl. Acids Res. 6:1081–1096.

[65] Tillo, D., Hughes, T. R. (2009) G+C content dominates intrinsic nucleosome occupancy. BMC Bioinformatics 10:442, 1-13.

[66] Yu, M., Ren, B. (2017) The three-dimensional organization of mammalian genomes. Annual Review of Cell and Developmental Biology, 33(1):265–289.

[67] Ganapathi, M., Palumbo, M. J., Ansari, S. A., et al. (2011) Extensive role of the general regulatory factors, Abf1 and Rap1, in determining genome-wide chromatin structure in budding yeast. Nucl. Acids Res. 39:2032–2044.

[68] Haran, T. E., Mohanty, U. (2009) The unique structure of A-tracts and intrinsic DNA bending. Quarterly Reviews of Biophysics 42:41–81.

[69] Levine, M. T., Jones, C. D., Kern, A. D., Lindfors, H. A., Begun, D. J. (2006) Novel genes derived from noncoding DNA in Drosophila melanogaster are frequently X-linked and exhibit testis-biased expression. Proc. Natl. Acad. Sci. U. S. A. 103:9935–9939.

[70] Zhao, L., Saelao, P., Jones, C. D., Begun, D. J. (2014) Origin and spread of de novo genes in Drosophila melanogaster populations. Science, 343, 769–772.

[71] Felsenstein, J. (1981) Evolutionary trees from DNA sequences: A maximum likelihood approach. J. Mol. Evol. 17:368–376.

[72] Felsenstein, J. (1989) PHYLIP-Phylogeny Inference Package (Version 3.2). Cladistics 5:164–166.

