## Supplementary figures Fig S1-S8. for "The triplet codons across the watershed between the non-living and living matters"

### Supplementary information to “The universal 3-base periodicity in genomes reveals a relationship between the evolution of the genetic code and the diversification of life”

Dirson Jian Li

**Abstract**

Supplementary figures Fig S1-S8.

#### Virus

Fig S1: SARS-CoV-2

Fig S2: SARS coronavirus Tor2

#### Archaea

Fig S3: Haloarcula hispanica

#### Bacteria

Fig S4: Rickettsiales bacterium Ac37b

#### Eukarya

Fig S5: Tribolium castaneum chrLG9

Fig S6: Gallus gallus chrZ

Fig S7: Homo sapiens Chr 21

Fig S8: Homo sapiens Chr 22

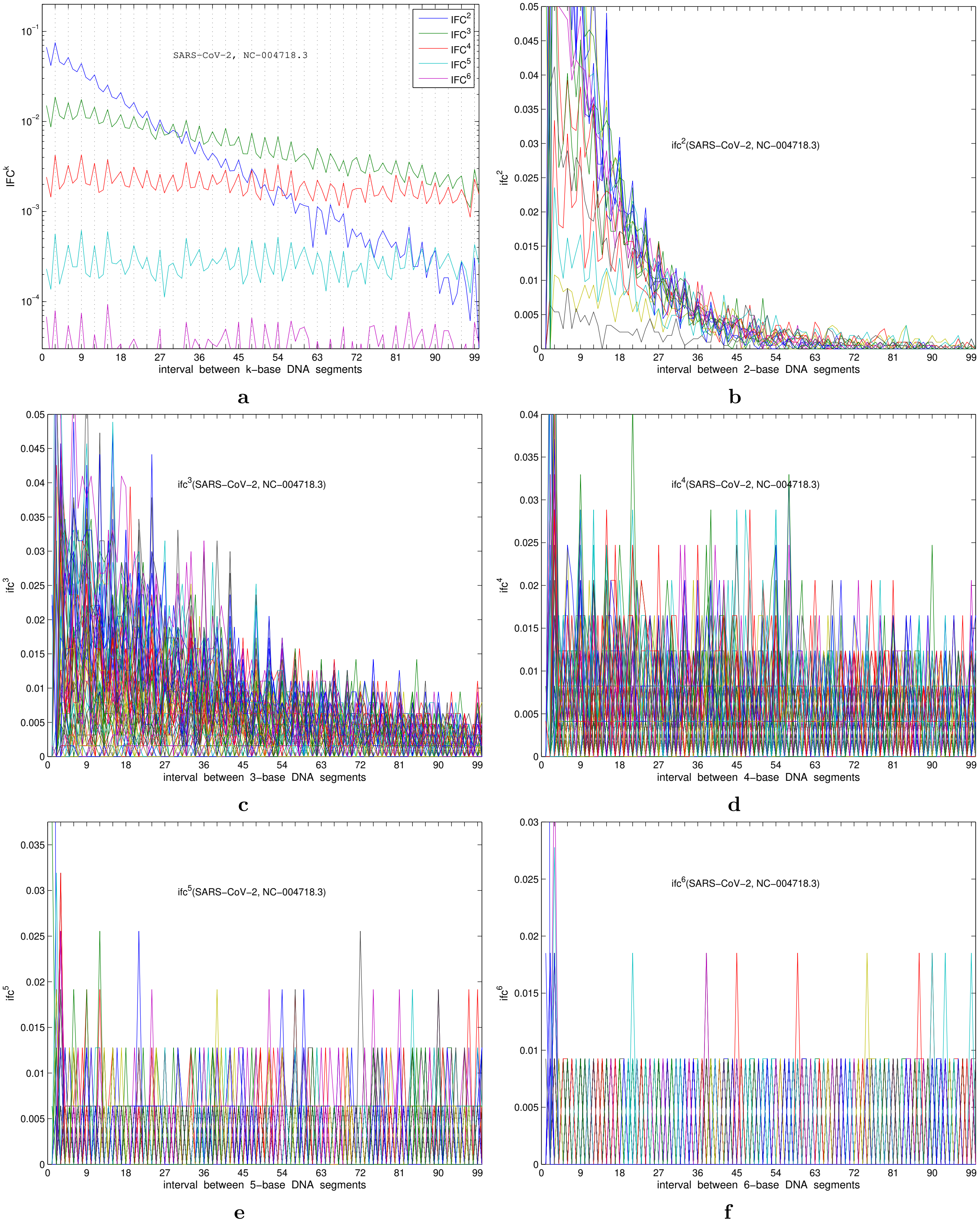


**Fig S1**: Interval fluctuation of codon $IFC^k$ and $ifc^k$ for SARS-CoV-2.

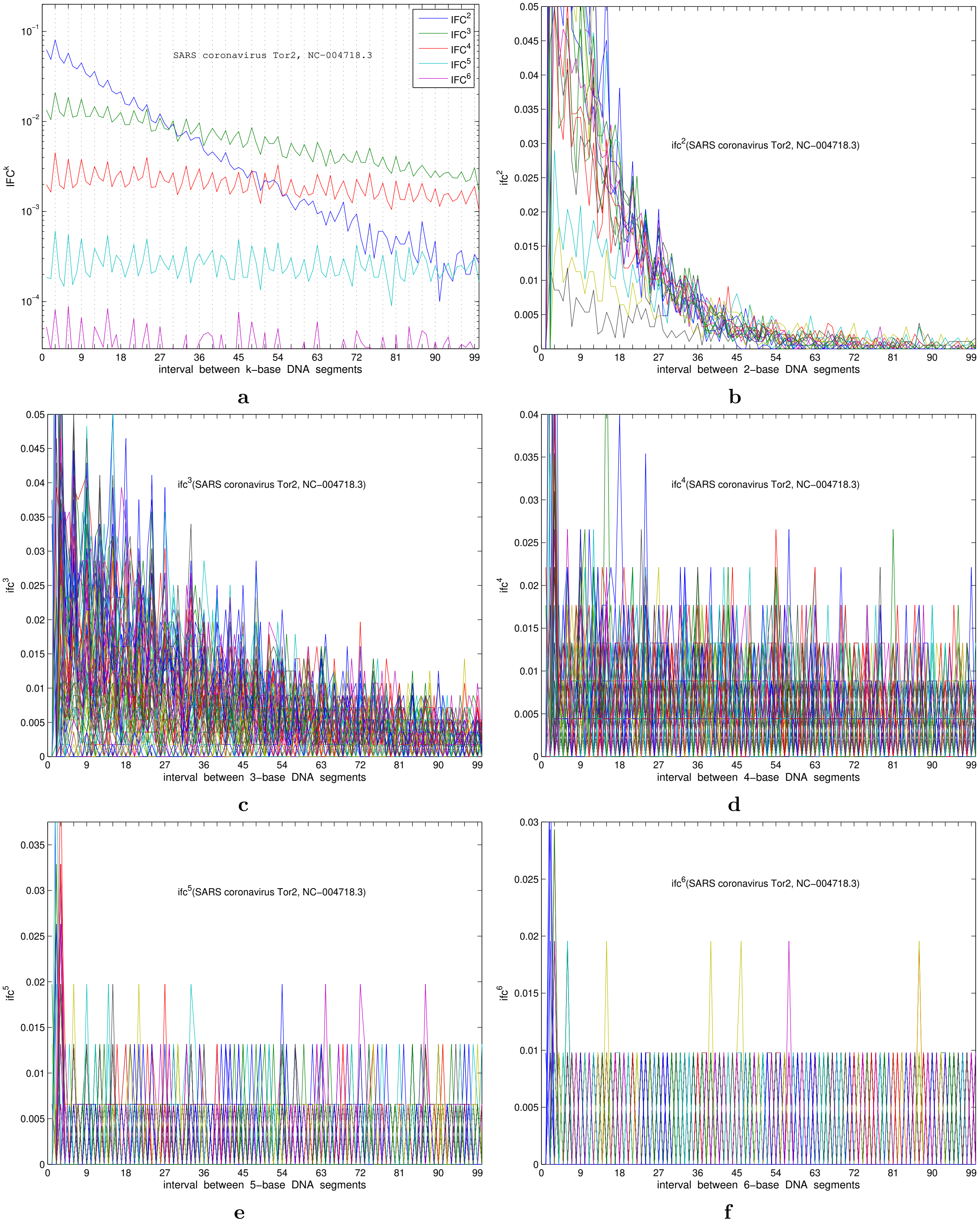


**Fig S2**: Interval fluctuation of codon $IFC^k$ and $ifc^k$ for SARS coronavirus Tor2.

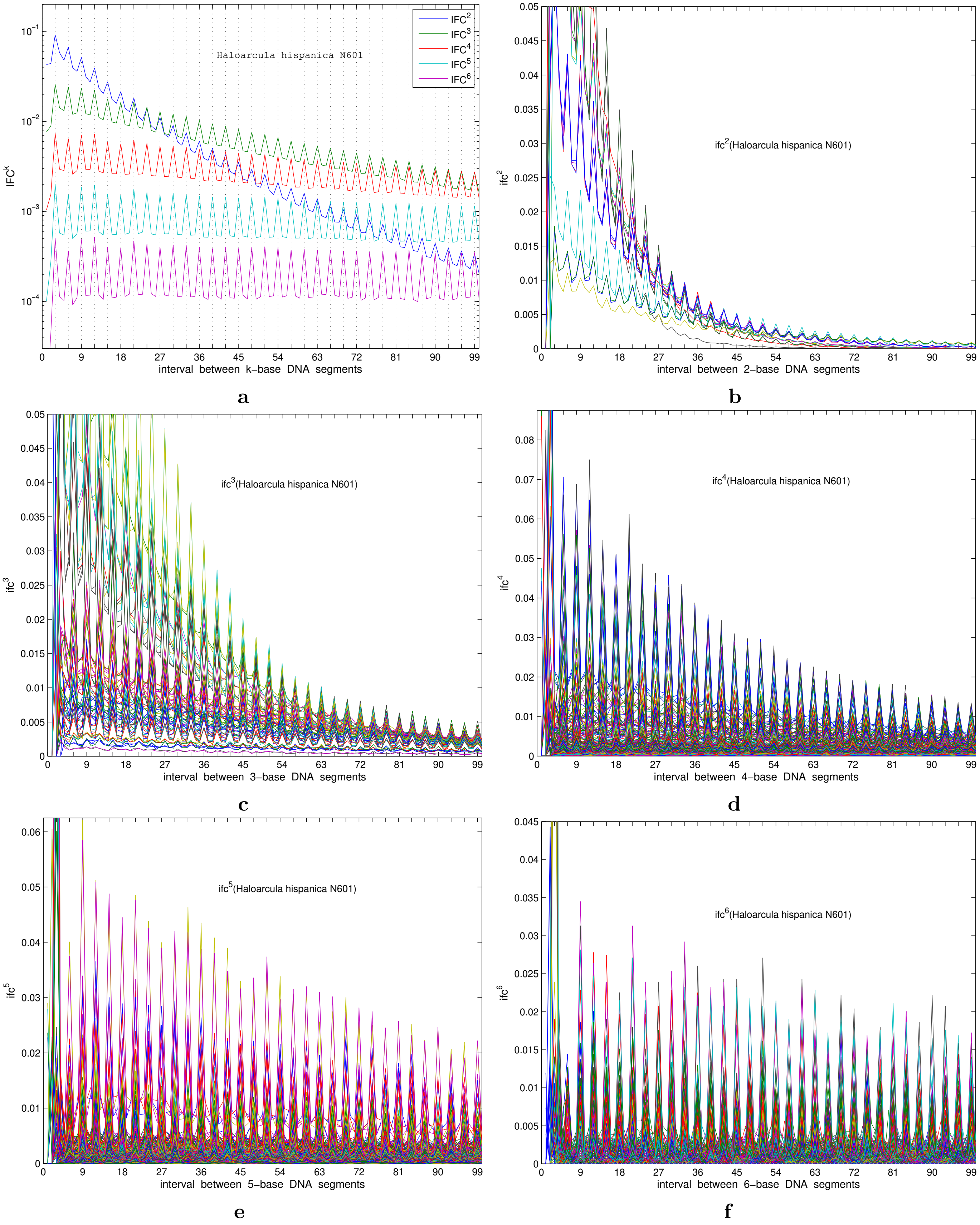


**Fig S3**: Interval fluctuation of codon $IFC^k$ and $ifc^k$ for Haloarcula hispanica.

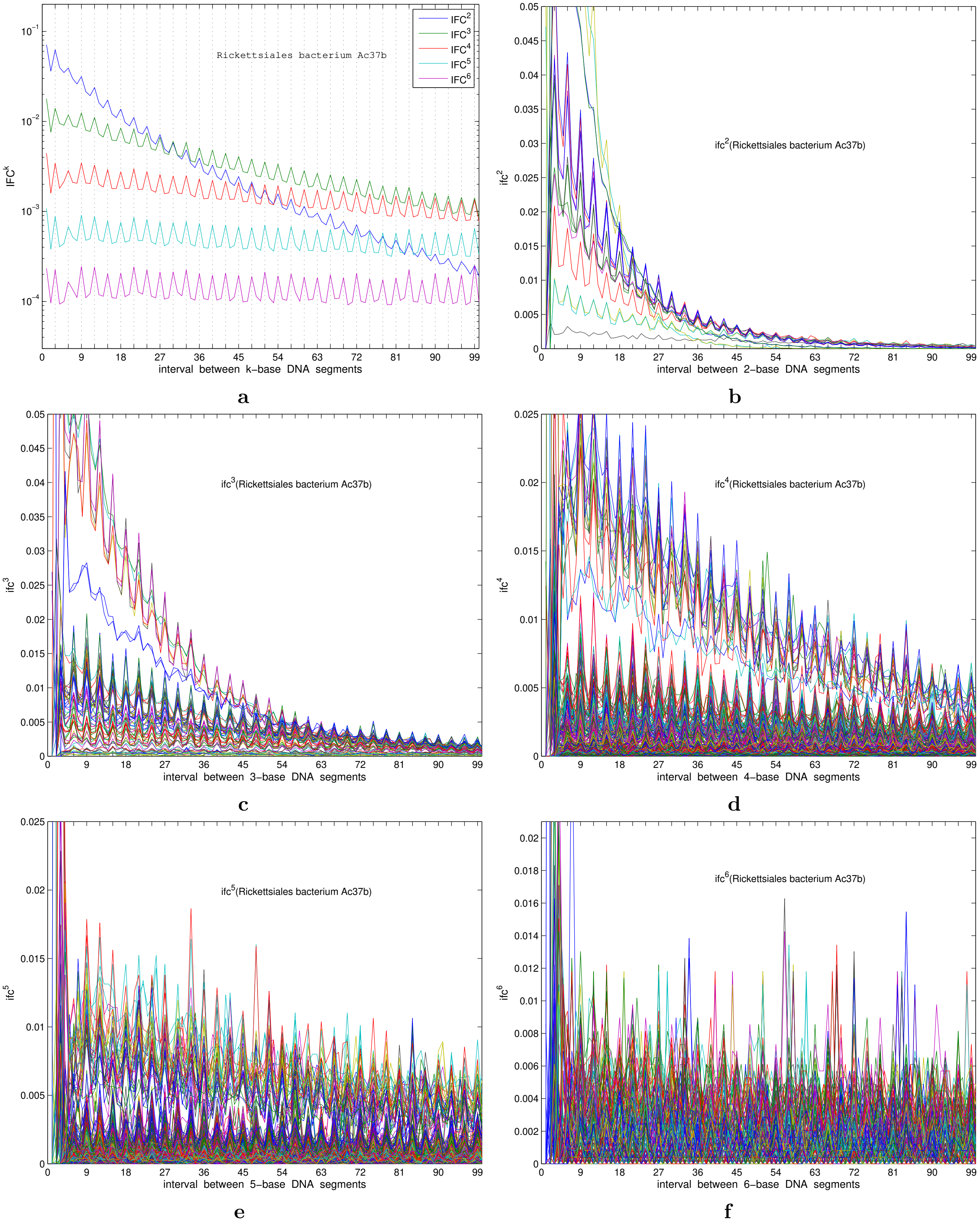


**Fig S**4: Interval fluctuation of codon $IFC^k$ and $ifc^k$ for Rickettsiales bacterium Ac37b.

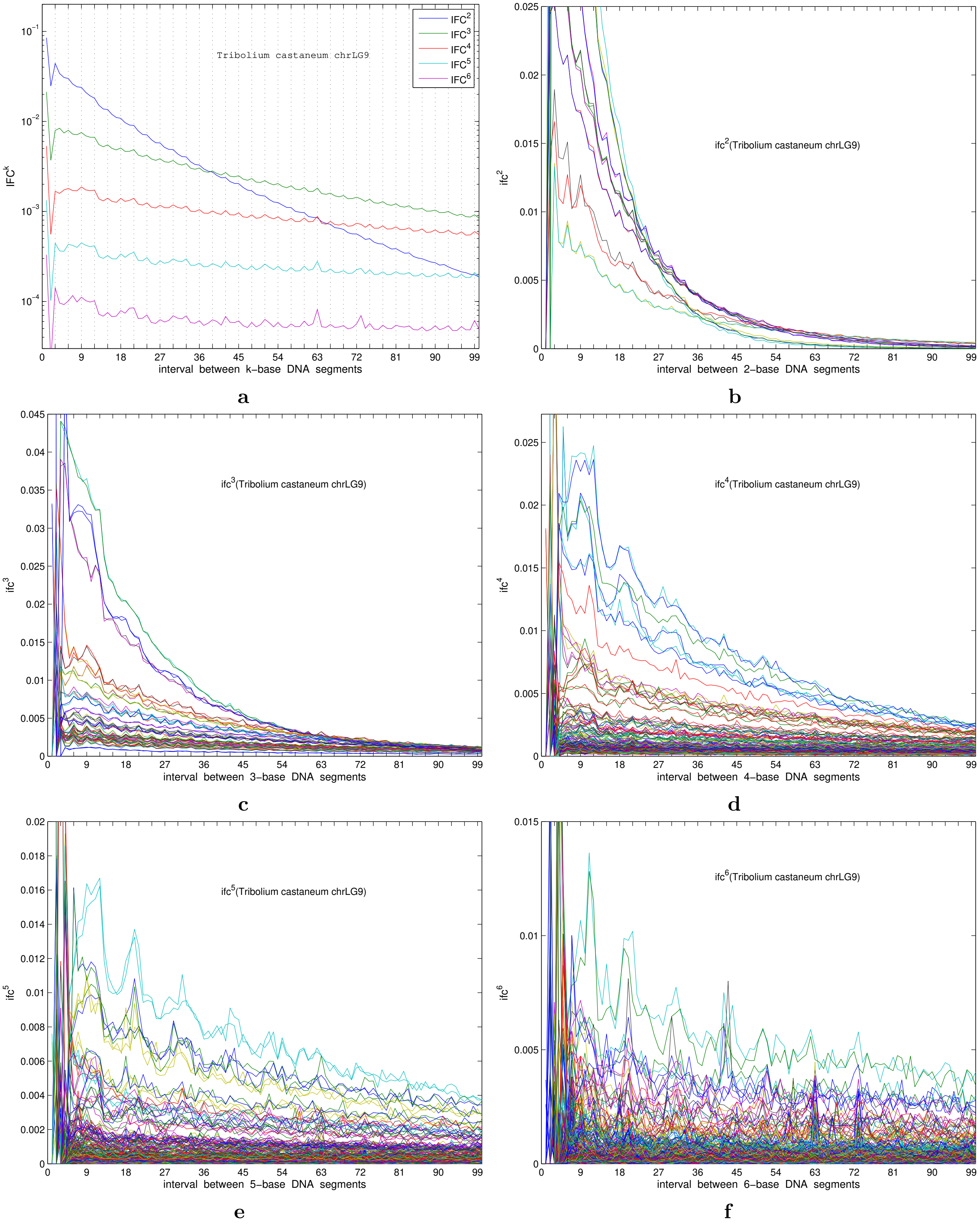


**Fig S5**: Interval fluctuation of codon $IFC^k$ and $ifc^k$ for Tribolium castaneum chrLG9.

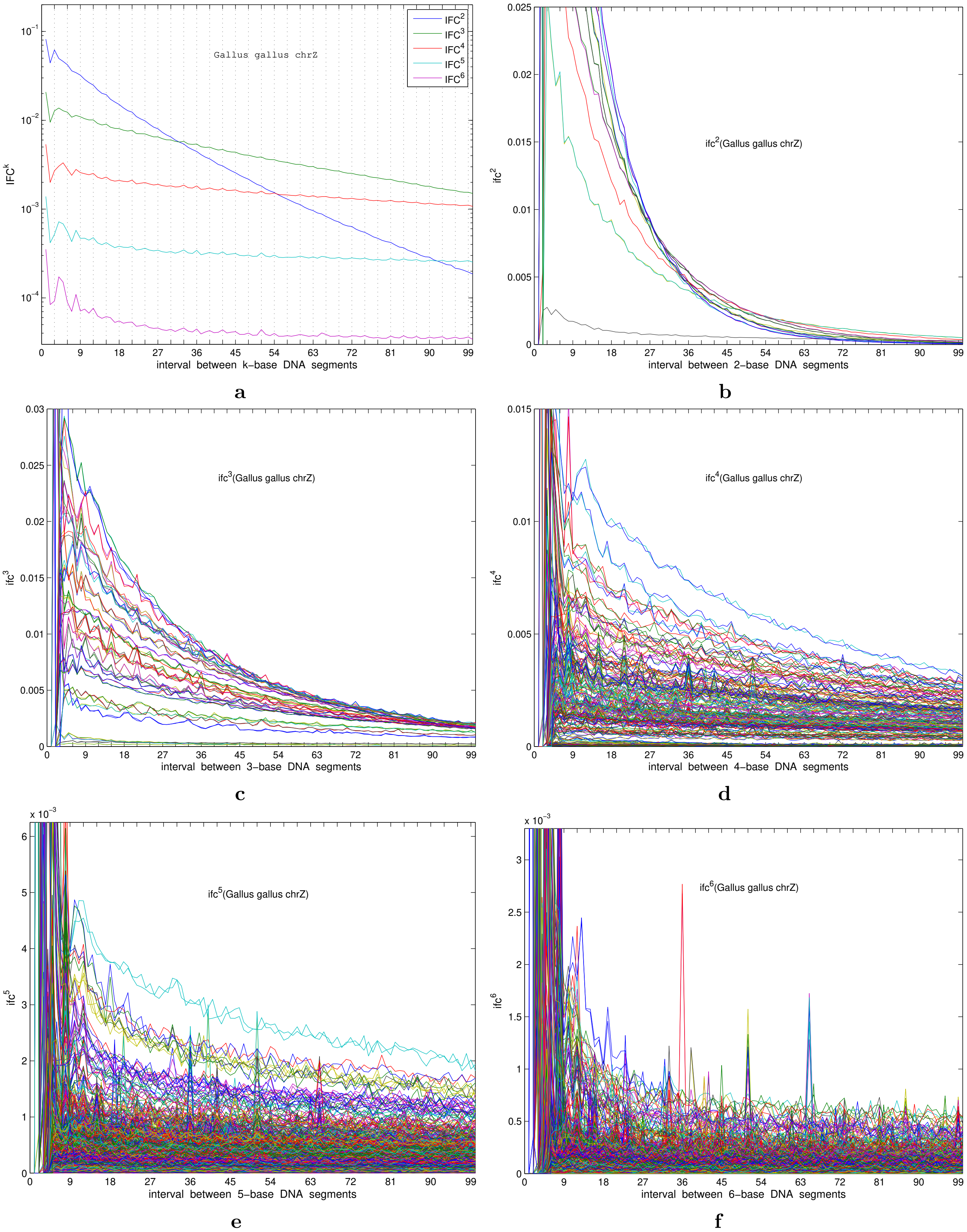


**Fig S6**: Interval fluctuation of codon $IFC^k$ and $ifc^k$ for Gallus gallus chrZ.

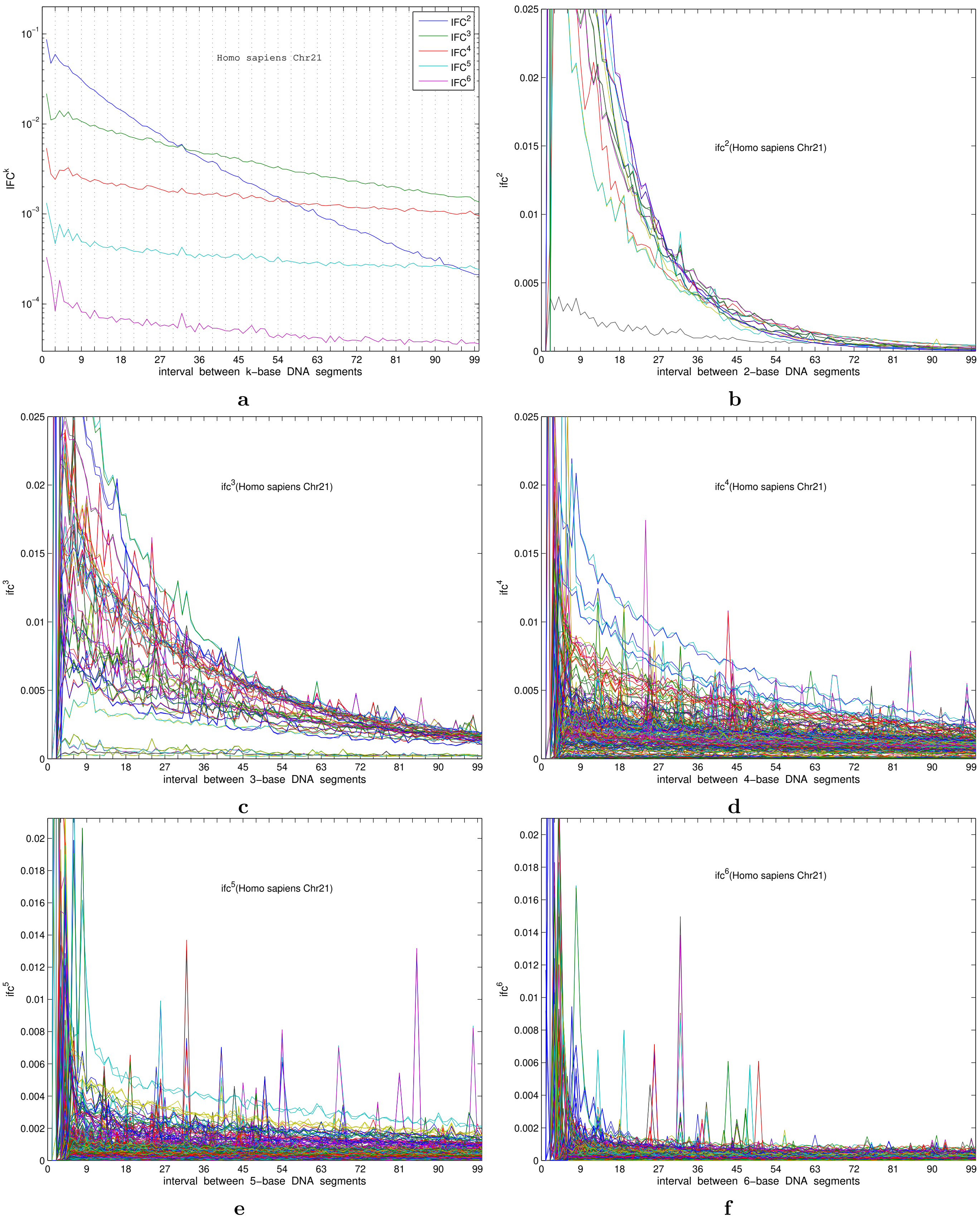


**Fig S7**: Interval fluctuation of codon $IFC^k$ and $ifc^k$ for Homo sapiens Chr 21.

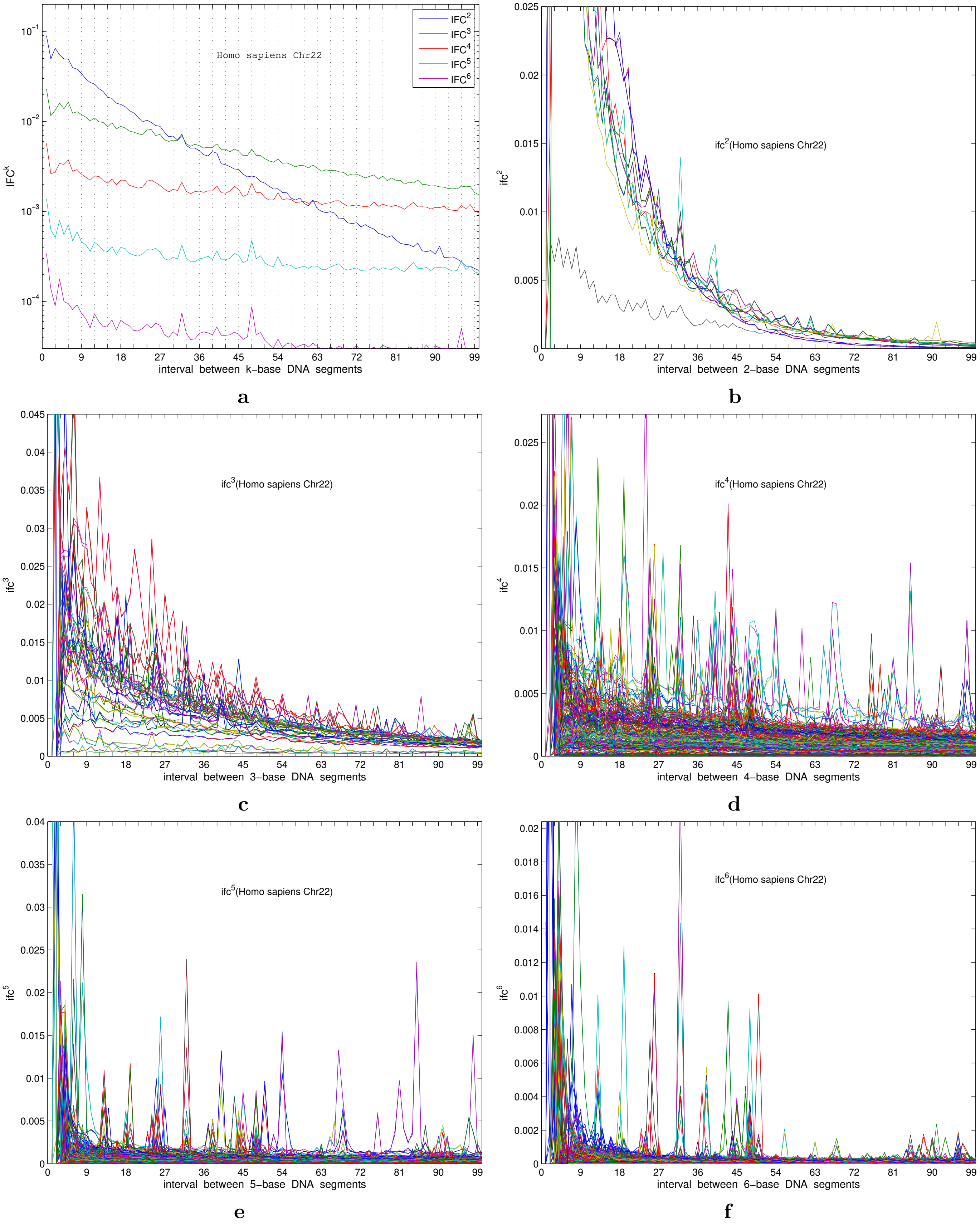


**Fig S**8: Interval fluctuation of codon $IFC^k$ and $ifc^k$ for Homo sapiens Chr 22.
